# Integrated time-lapse and single-cell transcription studies highlight the variable and dynamic nature of human hematopoietic cell fate commitment

**DOI:** 10.1101/101428

**Authors:** Alice Moussy, Jérémie Cosette, Romuald Parmentier, Cindy da Silva, Guillaume Corre, Angélique Richard, Olivier Gandrillon, Daniel Stockholm, András Páldi

## Abstract

Individual cells take lineage commitment decisions in a way that is not necessarily uniform. We address this issue by characterizing transcriptional changes in cord blood derived CD34+ cells at the single-cell level and integrating data with cell division history and morphological changes determined by time-lapse microscopy. We show, that major transcriptional changes leading to a multilineage-primed gene expression state occur very rapidly during the first cell cycle. One of the two stable lineage-primed patterns emerges gradually in each cell with variable timing. Some cells reach a stable morphology and molecular phenotype by the end of the first cell cycle and transmit it clonally. Others fluctuate between the two phenotypes over several cell cycles. Our analysis highlights the dynamic nature and variable timing of cell fate commitment in hematopoietic cells, links the gene expression pattern to cell morphology and identifies a new category of cells with fluctuating phenotypic characteristics, demonstrating the complexity of the fate decision process, away from a simple binary switch between two options as it is usually envisioned.

## Introduction

Hematopoietic stem and progenitor cells (HSPC) give rise to all the cellular components of blood. The major stages of differentiation and the key genes participating in this process are now well characterized [1]. According to the classical view, haematopoiesis is a hierarchically organised process of successive fate commitments, where differentiation potential is progressively restricted in an orderly way over cell divisions. There are several variants of the model [2-6]. In all cases the first fate decision is a binary choice taken by multipotent progenitors (MPP), which leads to two different committed progenitors (for the purpose of simplicity, these progenitors are designed here as common myeloid (CMP) and common lymphoid progenitors (CLP)). In molecular terms, the choice is believed to be the result of the strictly regulated activation of master regulator genes and their underlying transcriptional network [7]. However, the strict hierarchical logic of classical models has recently been challenged by a number of *in vivo* and *in vitro* studies [8,9]. Single cell gene expression studies have revealed a much higher heterogeneity of cell subtypes that can be detected using a combination of surface markers [10]. It is not surprising that the number of identifiable cell types increases with the resolution of the detection method. Although correct cell type classification is a key step in understanding the cell fate decision issue, it cannot reveal the dynamic features of the fate commitment process and leaves a number of unanswered questions. Do different phenotypic forms represent different cell types or different stages of the same process? How does the transition between the forms occur? How long does it take?

Until recently, fully deterministic explanations were predominant, but recent studies have suggested other alternatives. Two different possibilities have been put forward. According to the first, the commitment process starts with the sporadic, independent activation of genes within the same cell. The simultaneous stochastic expression of regulatory genes specifying different lineages creates a multiprimed intermediate state that enables these cells to choose one of the lineages [11-15]. A coherent lineage-specific expression profile would then emerge from this metastable state. According to the second, commitment is preceded by transcriptome fluctuations between different lineage-biased states [16-18]. Surprisingly the time scale of transformations related to the cellular fate decision process remains largely unexplored. The transcriptome of the same cell can be analysed only once, because the cell is destroyed by RNA extraction. Therefore, indirect approaches are required to identify trends and patterns in time series.

We addressed the issue of the dynamics and the time-scale of the commitment process by integrating single-cell quantitative RT-PCR, cell division history and morphological changes determined by time-lapse analysis. Contrary to the common strategy consisting in isolating defined cell subpopulations with the help of specific surface markers and characterizing their gene expression profiles at the single-cell level [19], we used an alternative approach. Individual cells were randomly isolated from the heterogeneous cord blood CD34+ cell fraction at different time points after cytokine stimulation and their gene expression profiles were determined using single-cell quantitative RT-PCR. The data provided a series of snapshots, showing the actual statistical distribution of single-cell gene expression patterns across the whole population. The structure of the population at the successive time points was revealed by unsupervised classification of the expression profiles according to their similarity using multiparametric approaches. The progression of the fate commitment process was deduced from the evolution of the population structure. At the same time, using time-lapse microscopy we tracked randomly isolated individual CD34+ cells and their progeny for several days after cytokine stimulation. We recorded the division history and the morphological changes of each cell in the clones. The population structure was deduced on the basis of the statistical analysis of these observations. The efficiency of the time-lapse approach in investigating cell fate decisions has been recently shown [20]. To reinforce this approach, the time-lapse and gene expression data were integrated into a coherent scenario. This was done by using CD133 protein expression levels to isolate cells with one or the other transcription profiles and record directly their dynamic phenotype, thereby providing a direct link between dynamic phenotype and transcription profile.

Altogether, our results revealed that fate decision is a dynamic, fluctuating process that is more complex than a simple binary switch between two options as it is usually envisioned.

## Results

### Single cell gene expression

The transcriptional profile of individual cord-blood CD34+ cells was determined at 0, 24, 48 and 72 hours after the beginning of cytokine stimulation (Fig.1A). Single-cell quantitative RT-PCR was used to quantify the mRNA levels of 90 different genes. A set of 32 genes was selected for their known function in the early differentiation of hematopoietic cells and were expected to inform on the functional state of the cells (see Supplemental Table 1.). A second set of 54 genes was chosen randomly from a list of genes known to be expressed in the hematopoietic lineage [21,22]. These genes provided an assessment of the overall transcriptional activity of the genome. Five additional genes were added to the list for their role in maintaining the pluripotent state in embryonic stem cells. A heat-map of all data and a violin plot of the expression profile of each gene at the four separate time points are shown in the supplemental information section (Fig S1). The normalized single-cell quantitative gene expression data obtained for the different time points were merged in a single database and screened for subpopulations by k-means clustering. The number of statistically distinguishable groups was inferred using gap statistics [23]. The groups were visualized on heat maps and on a two-dimensional plot using t-stochastic neighbour embedding (tSNE) [24]. Although every cell had a unique gene expression pattern, this approach enabled us to clearly identify sub-groups of cells in the population on the basis of the statistical similarity of their gene expression patterns (Fig 1B).

**Fig.1.**
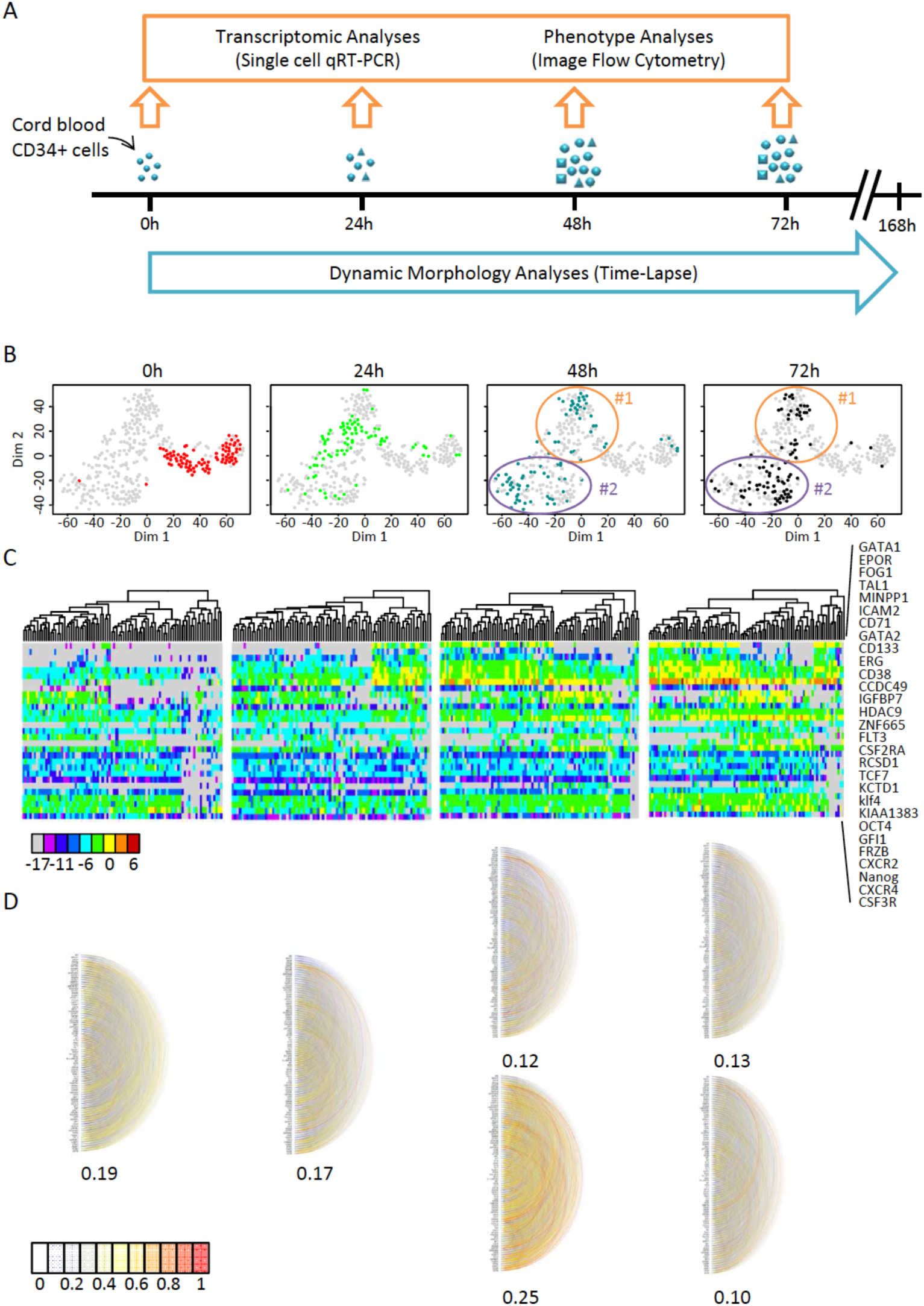
Transcriptional profile of cord blood-derived CD34+ cells at t=0h, t=24h, t=48h and t=72h after the beginning of the experiment. A. CD34+ cells were isolated from human cord blood and cultured in serum-free medium with early-acting cytokines. Single-cell qRT-PCR was used to analyze single-cell transcription at 0h, 24h, 48h and 72h. At the same time, individual clones were continuously monitored using time-lapse microscopy. B. t-SNE map of single-cell transcription data. The four panels show analysis of the same data set with each point representing a single cell highlighted by different color depending on the given time point. The two clusters identified by gap statistics at t=48h and t=72h are surrounded by an ellipse and numbered #1 and #2 for multipotent- and CMP-like cells. Note the rapid evolution of the expression profiles. C. A heat-map representation of the expression levels of a subset of genes that strongly contributed to the differentiation of the different groups (as detected by PCA; see Fig.S2) and cluster analysis of expression profiles at the different time points show the rapid evolution of gene expression. The list of the genes used (shown on the right) includes well-known genes acting during hematopoietic differentiation but also many randomly selected genes. The color code for expression levels is indicated below. D. Pairwise correlations between every gene analyzed using single-cell quantitative RT-PCR. Each semicircle represents the correlation between the two genes. The strength of the correlation is indicated by the color code. The average of all correlation coefficients is given separately for each group. The two clusters identified on t=48h and t=72h are represented separately. Note the transient increase of the average correlation in cluster #2 at t=48h.

Non-stimulated CD34+ cells isolated from cord blood represented the t=0h time point. A heat-map of the single-cell transcriptional profiles of genes contributing significantly to the identification of subgroups (Fig S2) showed that this population of cells was heterogeneous. Several genes reported to play a role in self-renewal, quiescence and other stem cell functions (CD71, CD133, CXCR4, GATA2 and FLT3) were expressed sporadically and at variable levels in a fraction of cells. Genuine pluripotent stem cell genes were also expressed at low level in a fraction of cells (Nanog, OCT4, KLF4). Nevertheless, no correlation was found between these genes (Fig.1D) and the statistical analysis did not reveal distinguishable expression patterns that could define cell types. The only detectable differences were donor-associated and probably reflected differences related to the processing of individual blood samples. Donor-specific differences disappeared at later stages.

The gene expression profile 24 hours after the onset of cytokine stimulation was found to be fundamentally different to t=0h cells. Almost every cell responded to cytokine stimulation by increasing transcript levels and generating a unique gene expression pattern (Fig 1C). When represented on the two-dimensional tSNE (Fig 1B) and PCA plots (Fig.S2A) the cells formed a single but dispersed cluster, well separated from the t=0h cells. In a fraction of cells moderate to high transcription of previously non-expressed hematopoietic regulator genes was observed in addition to that already seen at t=0h. For example, the expression of GATA1, GATA2, PU1, CD71, FOG1, CD133 or EPOR increased or was more frequent than at t=0h. In some cells all these genes were expressed simultaneously. Nevertheless, no distinct subpopulations could be identified at the resolution of our approach. The pairwise correlation coefficients between genes remained low (Fig 1D). Strong correlations were only between housekeeping genes (for example, GAPDH with ACTB). It is therefore likely that the patterns observed at 24 hours resulted from essentially uncoordinated up-regulation of gene transcription and led to a highly heterogeneous cell population. This is a transition state reminiscent of the reported multi-lineage primed state with simultaneous expression of lineage-affiliated genes specifying alternative cell fates [11,14].

The first signs of coordinated differential gene expression appeared at t=48h after cytokine stimulation. At this stage, two distinct gene expression patterns emerged from the highly variable background of earlier stages. The two clusters are clearly distinguishable on the tSNE plot (Fig.1B) and identified by gap statistics. They are also easily seen on the heat-map representing gene expression levels (Fig1C). Cluster#2 comprised cells with simultaneous expression of genes characteristic of erythro-myeloid progenitors (CMP) such as GATA1 and EPOR [7]. The expression profile of the cells in cluster#1 was characterized by the strong expression of genes reported for multipotent cells, CD133, GFI1, KLF4 or FLT3 and the lack of expression of GATA1 and EPOR. Although this pattern is reminiscent of a self-renewing, multipotent profile, it is difficult to determine the exact identity of these cells at the level of resolution used in our study [25]. Typical genes for pluripotent stem cells NANOG and OCT4 were expressed at moderate level in many cells from both clusters (Fig.1C and Fig.S1). Randomly selected genes were also good predictors for the two groups of cells. Only a small fraction of cells could not be classified in one of the two main clusters at t=48h (Fig.1B). The tendency observed at t=48h was further reinforced by t=72h. The cells in cluster #2 with CMP-like profile represented more than the half of all cells (Fig.1B-C). We observed a strong but transient increase in the number of highly correlated genes in this group (Fig.1D) indicating a state transition with the coordinated increase of the expression of some genes leading to a committed cell state. Indirectly, this suggested that the cluster#1 profile was more in continuity with the previous profile observed at t=24h and that the cluster#2 profile at t=48h represented a transition to a new pattern.

Taken together, these single-cell gene expression observations revealed that the cell-fate decision process in cytokine stimulated CD34+ cord blood cells occurred during the first two days. Initially, each cell responds to cytokine stimulation by an uncoordinated change in gene expression, which is followed by the emergence of two distinct gene expression patterns reminiscent of the two known major types of hematopoietic progenitor cells. Although indications of this second change may appear as early as 24 hours after stimulation, the two distinct gene expression patterns are clearly distinguishable at 48 hours and consolidated by 72 hours. By this stage almost every cell seems to have adopted one profile or the other.

### Time-lapse tracking studies

In order to integrate the gene expression snapshots into a dynamic scenario, we made time-lapse records of individual CD34+ cells under *in vitro* conditions identical to those in the single-cell gene expression studies. We imaged individual cells in microwells at a rate of 60 frames per hour for 7 days (Fig.2A). Using a semi-automatic image analysis approach we established individual clonal pedigrees, and recorded cell cycle durations and major morphological changes. As shown in Fig.2B,C, the pedigrees of individual clones were highly variable but shared some general features. Some clones produced only a few cells during the observation period, while others proliferated faster and produced up to 30-40 siblings. We focused our attention on the first three generations. As reported for cells cultured in early acting cytokines [26], the first cell cycle was exceptionally long in all clones. The division of the founder cell occurred between 35 to 80 hours after the start of culturing with the median cell cycle length being 58 hours (Fig.2). We questioned whether the culturing of isolated cells in microwells where direct contact with the other cells was not possible - influenced cell cycle length. To measure the division rate in a population context, the cells cultured together were labelled using Cell Trace Violet. The results (Fig.S3) showed that the cells had similar division profile regardless of whether they were cultured individually or in population. The unusually long first cell cycle was particularly important when interpreting results. It implied that the transition from the initial to the multi-lineage primed transcription profile followed by one of the two types of progenitor-like profiles observed at 24 and 48 hours after CD34+ cell stimulation occurred during the life of the founder cell, before the first mitosis.

**Fig.2.**
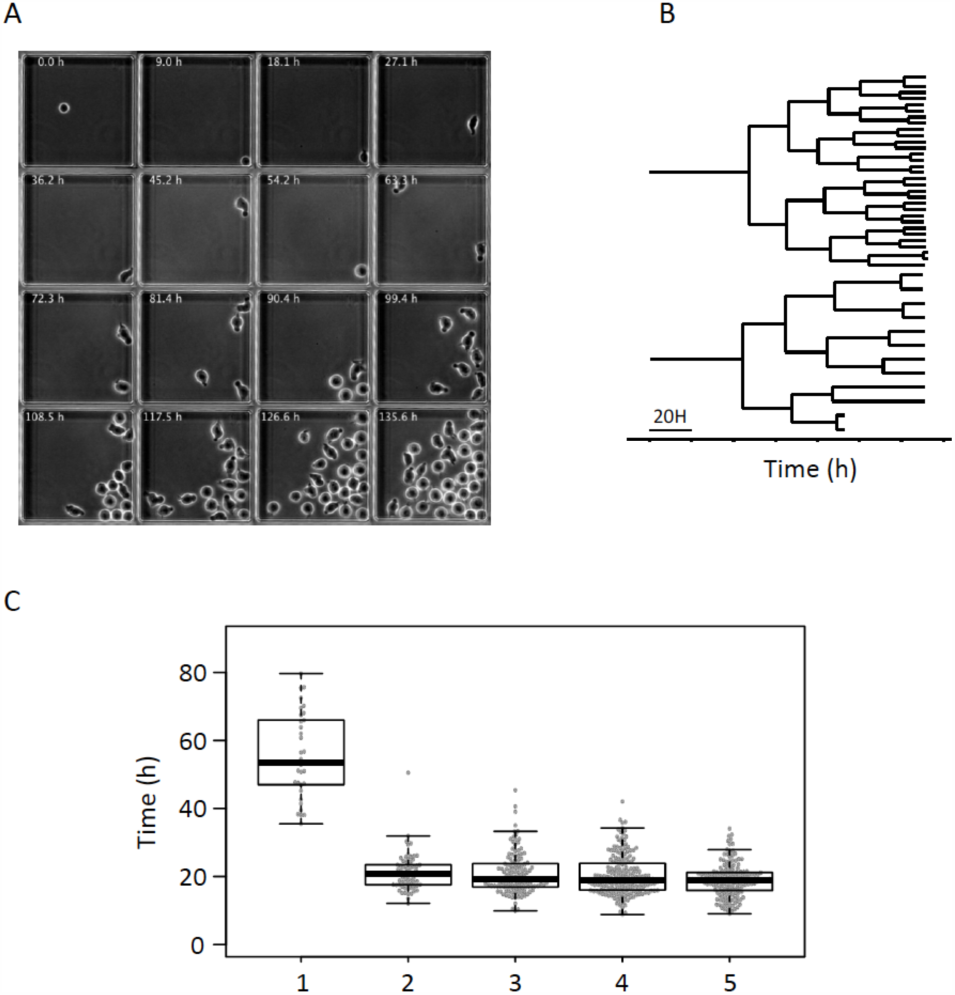
Time-lapse tracking of individual clones. A These frames extracted from a representative time-lapse record, show a microwell containing a single founder cell, which divides to produce a clone. Each individual cell was tracked and their morphological characteristics were recorded. B Two representative lineage pedigrees obtained from the time-lapse record. The strong difference in clone size observed at the end of the record is established gradually after the third cell division. C. Box-plot representation of cell cycle lengths obtained from the time-lapse records of every clone. Note the long first cell cycle. Subsequent cell cycles have comparable lengths with a slight tendency to become shorter.

Previous studies have demonstrated that there is a connection between cell morphology and the differentiation potential of CD34+ cells. Two major morphological forms have been described in the CD34+ cord blood cell fraction. Polarized cells are capable of active motion with the help of lamellipodia and possess, on their opposite end, large protrusions called uropods. These cells have been found to retain primitive self-renewing and stem cell functions [27,28]. The second morphological type is round. These cells have been considered as already engaged in differentiation [27,28].

Time-lapse records revealed that the two cell morphologies were not permanent; most cells were able to switch between forms several times during the cell cycle. After recovering from the stress of isolation and manipulation, founder cells acquired polarized morphologies within a few hours, developing uropod and starting to move actively (see Movies 1 to 3 in the Supplemental information section). During the first cell cycle, cells mostly conserved the polarized form and switches between the two morphologies were rare. As indicated above, the first cell division occurred on average at 58 hours and the average lengths of subsequent cell cycles were around 20 to 22 hours. The daughter (second generation) and granddaughter (third generation) cells were able to switch between the two morphologies at much higher frequency compared to the founder cells. In order to quantify these events, we manually tracked each cell and recorded each switch. Representative profiles are presented on Fig.S4.

In order to compare quantitatively the dynamic phenotype of cells we calculated three parameters based on their dynamic profile. The first parameter was calculated as a ratio of the time a given cell spent as a round shape compared to the time spent as a polarized shape. This parameter was close to 0 for stable polarized cells, and 1 for stable round cells. Intermediate values correspond to the fraction of time cells spent in round shape. The second parameter was the frequency of morphological switches during the cell cycle. This parameter expressed the cell's ability to maintain a stable morphology. The third parameter was the cell cycle length. When cells were represented as individual points in the space determined by the three parameters, we identified three major categories (Fig.3A). The first category included cells with mainly polarized shapes; the second category was composed of cells with predominantly round shapes. The cells in the third category switched shape frequently, generally fluctuating between both morphologies (Fig.3A)

**Fig.3.**
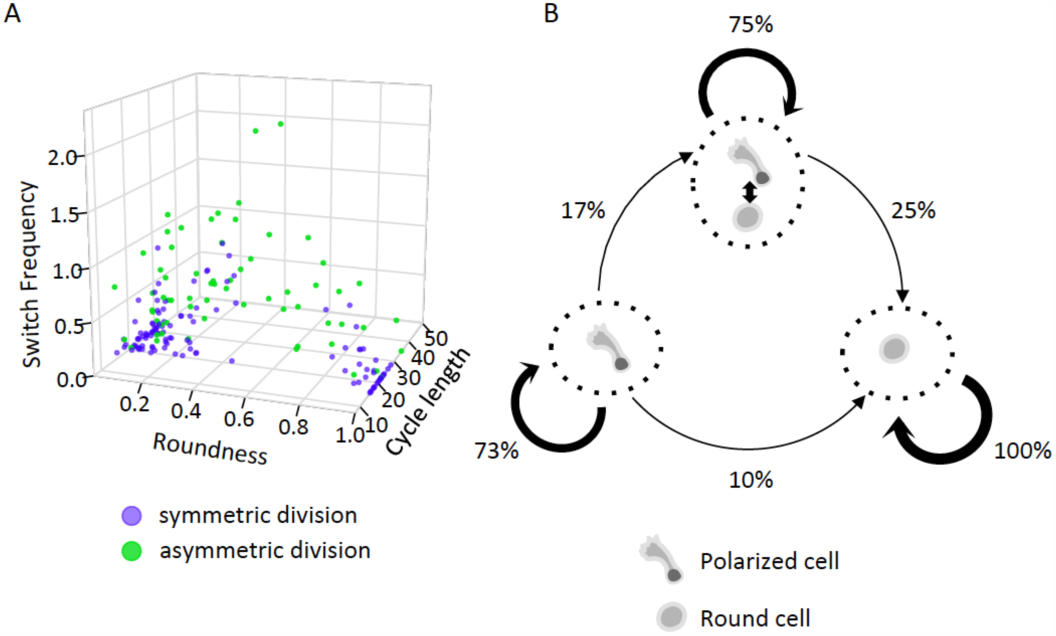
Quantitative analysis of dynamic phenotypes as determined by time-lapse data. A. Association between the morphology, switch frequency, cell cycle length and the type of cell divisions of second and third generation cells. Each point represents a single cell. Siblings with different switch behavior and morphology (in green) are usually characterized by high switch frequencies and frequently derived from the asymmetric division of the maternal cell. Symmetric divisions more frequently resulted in daughter cells (in blue) with similar dynamic behavior and morphologies. Division symmetry/asymmetry in the context of this work refers to the dynamic phenotype of the cells. The morphology is given as a ratio of time spent in round/polarized shape by a cell during the cell cycle. Switch frequency is given in number of morphological transformations per hour. Cell cycle length is in hours. B. Dynamic phenotype change during the first two cell divisions as determined on the basis of time-lapse records. Three different dynamic phenotypes were identified: stable polarized, frequent switchers and stable round. Cells tended to transmit dynamic phenotypes to daughter cells during cell division. Polarized and frequent switchers produced round cells, frequent switchers always produced by polarized mothers. Phenotypic change is not associated with asymmetric division; it can occur at any time in the cell cycle. Since round cells always produce round daughters, the whole process is biased and the proportion of this phenotype increases.

When sister cell pairs were examined, it became obvious that many displayed very similar dynamic phenotypes. In some cases, periods of stable morphology and switching events coincided almost perfectly (Fig.S4). We considered these cells were produced by symmetric cell division. In other cases, division was asymmetric and the two sister cells behaved differently. In the most extreme cases, one sister cell adopted a stable round form and the other a stable polarized form immediately after division. However, most of the asymmetric divisions produced siblings with obviously different dynamic phenotypes. Based on the above-mentioned parameters, we used k-means clustering to classify cells derived from a symmetric or asymmetric division (Fig.3A). This classification confirmed that most asymmetric divisions produced cells with highly fluctuating dynamic phenotypes.

We calculated the frequency with which a cell with a given dynamic phenotype was produced by a mother cell with similar or dissimilar phenotype (Fig.3B). Maternal cells clearly tended to transmit the dynamic phenotype to daughter cells. We also observed regularity with which phenotype changes occurred in daughter cells. Polarized cells were systematically produced by polarized cells. At lower probabilities, both polarized and fluctuating cells could produce stable round phenotype cells. Round cells always gave rise to round siblings (Fig.3B). Following these simple rules, the cumulative outcome of the process was the gradual increase of round cells in the population. Cells with fluctuating morphologies appeared to be intermediate form between polarized and round cells. Since 25% of daughter cells conserved this phenotype, the fluctuating intermediate cells persisted in the population. On static snapshots however this category remained undetectable: only polarized and round cells were observed. A polarized form was considered to be a feature of multipotent cells and the round form a committed myeloid progenitor phenotype [27,28].

### Coupling the molecular and cellular scales

The dynamically fluctuating behaviour we have described here for the first time represent a transition between the two states and reflect a “hesitant” but incomplete fate determination process. Since we detected only two major transcription profiles but observed three different dynamic behaviours, it is possible that “hesitant” cells are not characterized by a clearly distinct transcription pattern. Morphology fluctuations may be accompanied by fluctuations in the transcript or protein levels of at least some key genes.

To test this assumption, we took advantage of the observation that the gene coding for the CD133 cell surface protein was expressed preferentially in one of the two transcription patterns detected at 48 hours (Fig.1B and C). Previous reports have established that CD133 protein is typically present in cells with polarized forms and accumulates in the uropod [27-29]. We confirmed this using image cytometry and immunohistochemistry on fixed cells (Fig.S5). Cells expressing high levels of CD133 were mostly polarized, while those with low levels of CD133 were round (Fig.S5). This observation explicitly established the link between the cell morphology and the transcription patterns detected by single-cell RT-PCR.

We used the CD133 protein as a proxy for the isolation of a cell fraction enriched either in polarized or round cells and recorded their dynamic phenotype. The “high” and “low/negative” CD133-expressing cell fractions were sorted 48 hours after cytokine stimulation, put in culture and tracked using time-lapse microscopy for an additional 48 hours. The cell-cycle length, switch frequency and division asymmetry were calculated. The “high CD133-expressing" cells and their progeny reproduced the three types of cells observed previously but in different proportions (Fig.4). Most of the cells displayed stable polarized morphologies or were frequent switchers; only a few cells displayed stable round morphologies (Fig.4). By contrast, the “low/negative" cells produced either stable round progeny or cells with fluctuating morphologies (Fig.4). The frequency of asymmetric division (in terms of dynamic behaviour) was higher in the “high CD133" population than in the “low CD133" population (78% *versus* 59%).

**Fig. 4.**
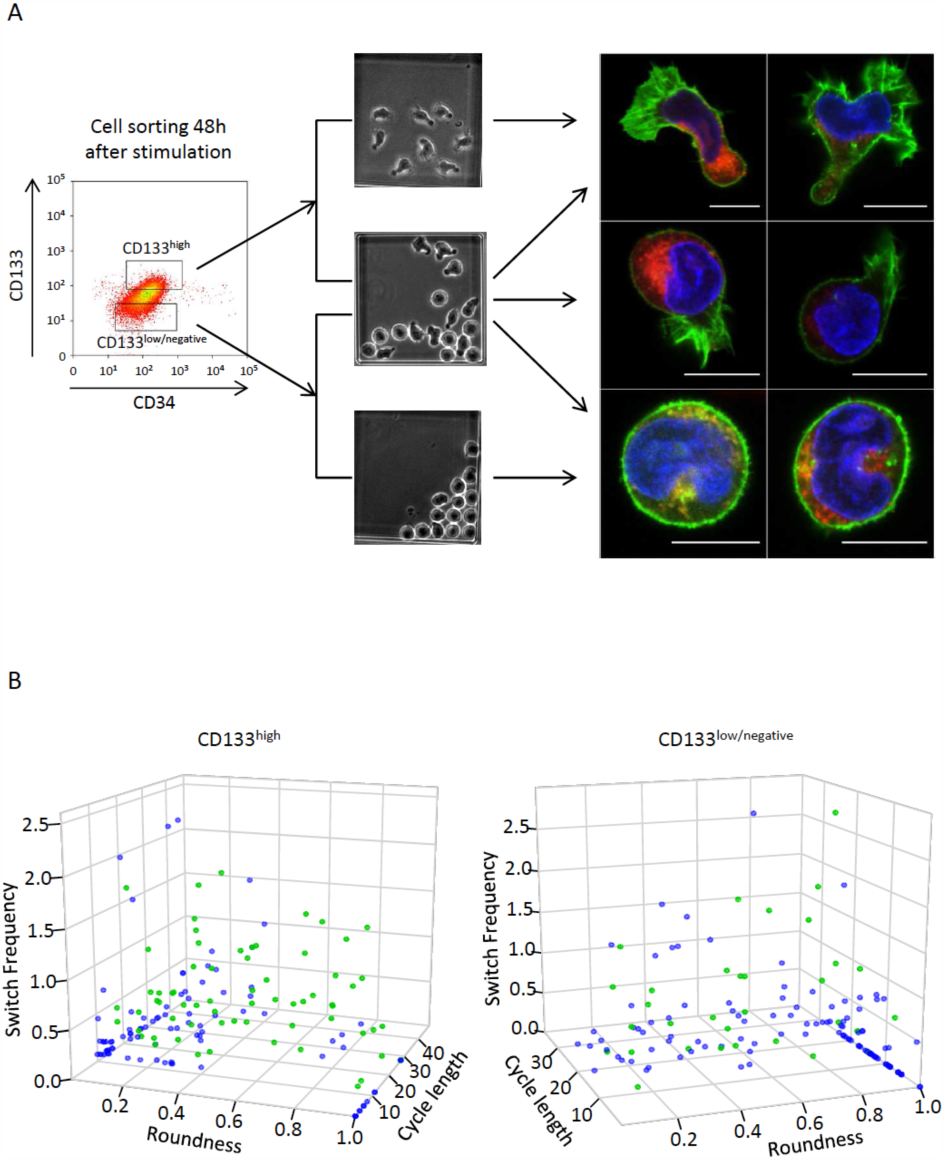
Isolation of polarized high CD133-expressing and round low CD133-expressing cells and time-lapse analysis of their dynamic phenotype. A. Cell sorting strategy to isolate cells with defined morphology on the basis of the CD133 surface protein level. The isolated cells were tracked by time-lapse microscopy. They produced cells with polarized, round or fluctuating dynamic phenotype (illustrated by the middle panel). Examples of cells with different morphology are shown on the right, as detected by confocal microscopy. Red: CD133 protein. Green: actin filaments detected by phalloidine. Blue: DNA. Note the preferential localisation of the CD133 protein in the uropode of polarized cells. Actin is concentrated in lammelipodia or evenly distributed in the periphery of round cells. B. Quantitative evaluation of the morphology, switch frequency and cell-cycle length of daughter and granddaughter cells derived from sorted “high CD133” (left panel) and “low CD133” (right panel) cells. Each point represents a single cell. The color code for symmetric and asymmetric division are the same as in Fig.3A. Note the higher frequency of frequent switchers and cells derived from asymmetric division of “high CD133” mothers.

These observations confirmed the idea that cells with stable round shape were derived from cells with polarized shapes and high CD133 levels, following a gradual transformation process that involved a transition period with fluctuating phenotype. The process of transformation did not correlate with the cell cycle; some cells reached a stable morphology rapidly while others fluctuated over several cycles. The process was accompanied by a gradual decrease in CD133 protein level in cells. We found no evidence that asymmetric divisions played a direct role in this process.

### Single-cell transcription profile of the multipotent stage

In order to determine which of the observed phenotypes correspond to the multipotent stage, we took advantage of recent observations demonstrating that the inhibition of histone deacetylase (HDAC) activity with a pharmacologic agent resulted in a substantial increase in their incidence in the cord blood CD34+ population [12,30,31]. We anticipated that this would increase the proportion of cells with transcription profile typical of the multipotential phenotype. Since valproic acid (VPA) was shown to be the most efficient [30], we used this agent to treat CD34+ cord blood cells stimulated by cytokines as above, before sampling transcription profiles. The increase of the CD90 marker, as analysed by flow cytometry confirmed that the VPA effect was already visible after 24 hours and gradually grew stronger during subsequent steps (Fig.5A and Fig.S6). The expression of CD34 and CD38 markers remained unchanged (Fig.S6). Although we did not analyse the *in vivo* potential of these cells, based on previous reports, we considered them enriched for *bona fide* multipotent cells. We performed single-cell RT-PCR 0, 24, 48 and 72 hours after the start of the experiment, as in control cells. At all four time points, cell populations were very heterogeneous. At each time point, the cells displayed a unique transcription profile (Fig.5B) and no identifiable transcription patterns appeared during the 72 hours of the experiment despite slight profile evolutions. Overall, transcription patterns in individual cells were reminiscent of the uncoordinated multi-lineage primed profile detected in control cells at 24 hours, but the two groups clustered separately on tSNE maps (Fig.5C). Since the cells did not divide during the first 48 hours, the increase observed in the multipotent cell fraction could not result from the selective proliferation of an initially small subpopulation of cells. Instead, this occurred because cells already present in the population changed the expression of many genes in response to the valproic acid.

**Fig.5.**
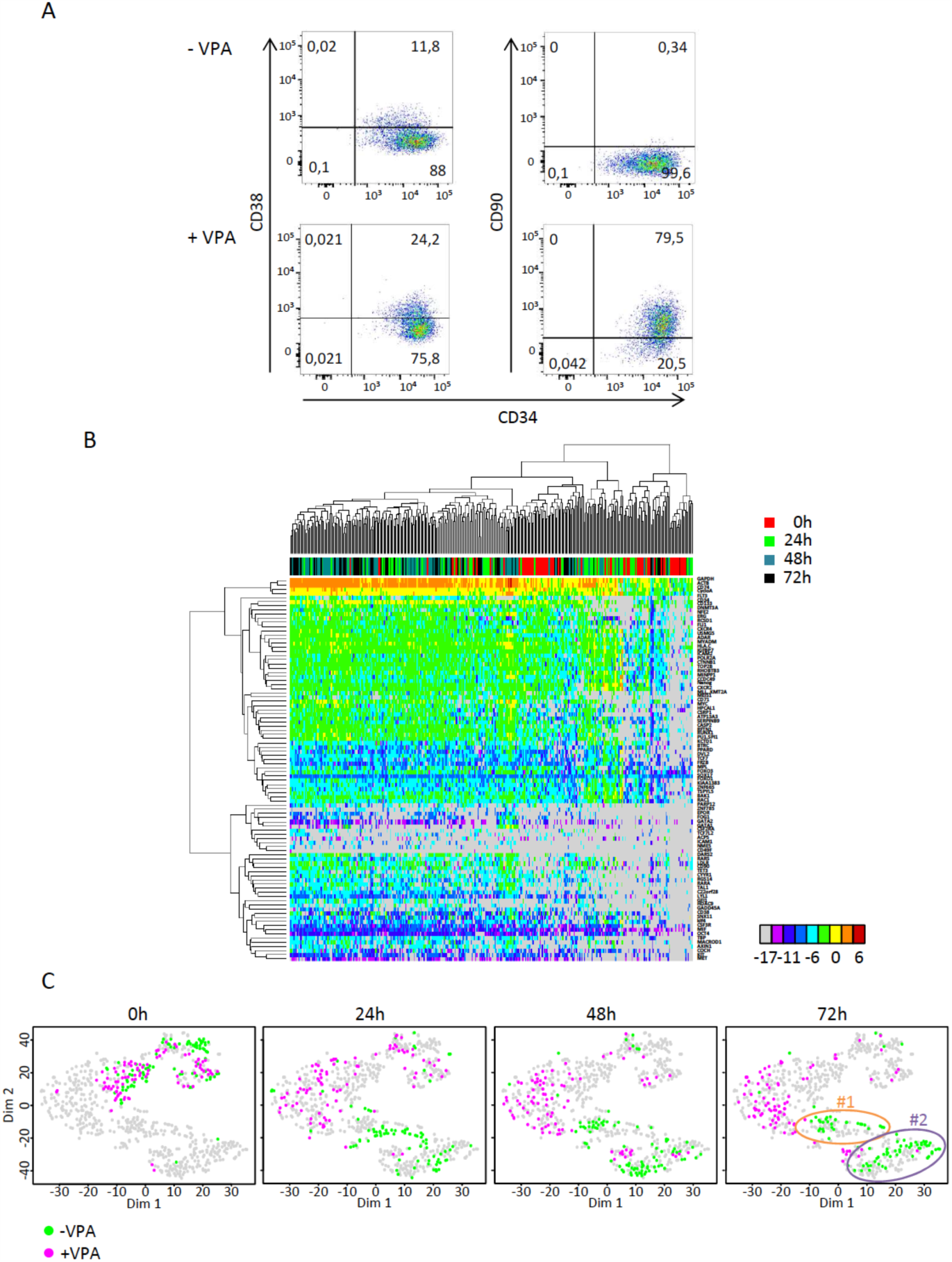
Transcriptional profile of cord blood-derived CD34+ cells treated with valproic acid (VPA) at t=0h, t=24h, t=48h and t=72h after the beginning of the experiment as compared to untreated normal control cells. A. A cytometric analysis of the effect of VPA on cord blood CD34+ cells shows increase in the CD90 protein in most cells while the CD34 and CD38 markers remain essentially unchanged. B. Heat-map representation of the expression levels of 90 genes as determined by single-cell qRT-PCR in VPA treated cells at t=0h, t=24h, t=48h and t=72h. The colour codes for the time points of cells are indicated on the right, the color code for expression levels are indicated below the heat-map. Note the high heterogeneity and lack of clear clustering of the expression patterns. C. tSNE plot representation of transcription data obtained for VPA treated-compared to untreated normal cell (data for these cells are the same as in Fig1.). The gene expression data obtained in the two experiments were mapped together. Each point represents a single cell and the cells at t=0h, t=24h, t=48 and t=72h are highlighted separately in the four panels. The color codes for VPA-treated (VPA+) and untreated (VPA-) are indicated below the panels. The clusters #1 and #2 identified at t=48 and t=72h in VPA- cells (see Fig1.) are indicated on the t=72h panel. Note the clear separation of the VPA+ and VPA- cells at every time-point except t=24h. Note also that VPA+ cells do not contribute to clusters #1 and #2 indicating that they do not acquire expression profiles typical of these cells.

## Discussion

In the present study we aimed to identify the initial stages of fate commitment in the CD34+ cell fraction of human cord blood and determine the typical time scale for these events. Without cytokine stimulation, CD34+ cells remain quiescent and die after a few days in culture. Early-acting cytokines allow these cells to survive, become metabolically active and enter the cell cycle [32] without showing overt signs of differentiation during the first few days. This creates ideal conditions for studying early events. Our experimental design combined continuous time-lapse observations with snapshots of high-resolution single-cell transcriptome analysis. The data can be integrated in a dynamic fate decision process scenario. Fate decision is necessarily accompanied by a change in the gene expression pattern. This is a multistep process. Firstly, upon stimulation, cells rapidly reach the multiprimed state, which is characterized by promiscuous gene expression pattern and predominantly polarized morphologies. This is an unstable phase and two distinct transcription profiles start to emerge before the end of the first cell cycle. The process by which cells relax from a multiprimed to more stable state is continuous and of variable length. Some cells reach stable morphology and coherent lineage-affiliated transcription profile by the end of the first cell cycle, which they transmit to daughter cells. Other cells divide into unstable daughter cells with dynamic “hesitant" behaviour. This behaviour is characterized by fluctuations between polarized, actively moving, amoeboid and round morphologies over several cell cycles, suggesting that instability can be transmitted mitotically. Although we have no formal evidence that the transcriptome of these cells also fluctuates, two observations suggest that this could be the case. Firstly, we only found two established transcription profiles that presumably correspond to polarized and round morphologies with high and low CD133 expression levels (Fig.1C). However, we observe three dynamic phenotypes, one of which is fluctuating. Secondly, cells isolated on the basis of high CD133 protein level were either polarized or fluctuating and low expressing cells were either round or fluctuating. This suggests that fluctuating cells may have intermediate levels of the CD133 protein and represent a transition between the stable polarized morphology and round morphology. This dynamic scenario reconciles assumption of a stochastic multi-lineage primed state with the idea of fluctuating transcriptomes [14,16] by suggesting that relaxation from the metastable multi-lineage primed state to the stable lineage committed state involves an uncertain “hesitant" phase of variable length.

Increased stochastic variation in gene expression may be responsible for the rapid shift away from the initial quiescent state and lead to the uncommitted multi-lineage primed state [11,14,15]. Cell division is not required for this process; it occurs during the first cell cycle following stimulation. Cells engaged on the path toward the new phenotype represent the committed state. The critical moment in this process is the transition between the two phenotypes when the old gene network has broken down but the new network is not yet assembled. We consider that cells with fluctuating morphologies represent this transition state. The rapidity of the transition may be dependent on the time required for the new gene expression network to settle to a stable state. Since phenotypic stability of a cell lineage largely depends on the frequency of transcription initiation and the stability of the resulting mRNA-s and proteins [33,34], the observed “hesitant” phenotype might be the consequence of rapid mRNA and protein turnover. The consolidation of the chromatin structure appears to be an essential element in this process, because, as shown in single cell transcription studies the HDAC inhibitor valproic acid delays the transition and blocks cells in a promiscuous gene expression pattern typical of multi-lineage primed state. Indeed, HDAC inhibitors have been shown to increase gene expression stochasticity by increasing chromatin acetylation [35].

In summary, in the current study we identified the earliest phases of fate commitment in human cord blood CD34+ cells and assigned a time scale to this process. We demonstrated that the rapid initiation of the process occurs within a single cell cycle and is followed by a dynamic transition state of variable length that may span several cell cycles. Since experimental conditions were constant, the changes observed are likely to reflect cell-intrinsic processes whereas the convergence toward a similar endpoint may reflect the constraints imposed by these conditions. From this perspective and in accordance with earlier theoretical and experimental work [36-40], fate decision appears to be a process of spontaneous variation/selective stabilisation reminiscent of trial-error learning, where each cell explores many different possibilities at its own rhythm by expressing a large variety of genes before finding a stable combination corresponding to the actual environment.

## Materials and methods

### Human sample and cell culture

Human cord blood (UCB) was collected from placentas and/or umbilical cords obtained from Etablissement Français du Sang (EFS), Saint Louis Hospital, France or from Centre Hospitalier Sud Francilien, Evry, France in accordance with international ethical principles and French national law (bioethics law n°2011-814) under declaration N° DC-201-1655 to the French Ministry of Research and Higher Studies. Human CD34+ cells were isolated from the mononuclear fraction of UCB samples using the autoMACSpro (Miltenyi Biotec, Paris, France) immunomagnetic cell separation system. They were then cryopreserved in Cryostor (StemCell, Paris, France) and stored in liquid nitrogen or used directly without freezing.

Cells were cultured at 37°C in a humidified atmosphere containing 5% CO2 in a 24-well plate in X-VIVO (Lonza) supplemented with 100 U/ml penicillin, 100 μg/ml streptomycin (Gibco, Thermo Scientific), 50 ng/ml h-FLT3, 25 ng/ml h-SCF, 25 ng/ml h-TPO, 10 ng/ml h-IL3 (Miltenyi Biotec, Paris, France) final concentration. Valproic acid (VPA) (Sigma Aldrich) was used at final concentration of 1,25mM.

### Single cell qRT-PCR

Single-cell qRT-PCR was carried out using the Fluidigm BioMark platform. DELTAgenes assays (life technologies) were used at a final concentration of 500nM for each of the 96 assays. Individual cells were sorted directly into a RT mix solution and spikes (life technologies) in a 96-well plate. RNA was denatured and reverse transcribed. of Premplification (20 cycles) of 96 specific cDNA was performed by denaturing the cDNA at 96°C for 5 secondes followed by annealing and extension at 60°C for 4 minutes. Unincorporated primers were removed by Exonuclease I and the preamplified products were diluted 5-fold. Amplification was performed with EvaGreen Supermix with Low ROX and inventoried DELTAGenes assays in 96.96 Dynamic Arrays on a Biomark System (Fluidigm). Ct values were calculated from the system’s software (BioMark Real-Time PCR Analysis, Fluidigm).

### Single cell Data Normalization

Ct values obtained from the Biomark System were normalized with the help of two externally added controls (spike 1 and spike 4, life technologies) according to a set of rules provided below. For each gene, inconsistent readings or “failed” quality control readings were removed. Cells with failed or inconsistent detection of spikes were removed. Expression values were calculated by subtracting the gene Ct value from the geometric average of Ct values from spike 1 and spike 4 in the corresponding cell. An arbitrary dCt value of −17 was assigned for all the genes with a dCt value of less than −17.

### Single cell qRT-PCR data analysis

Analyses of qRT-PCR single cell data were performed with R software (R Core Team (2012). R: A language and environment for statistical computing. R Foundation for Statistical Computing, Vienna, Austria. ISBN 3-900051-07-0, URL http://www.R-project.org/) using Heatmap3, factomineR, kmeans, ggplot2 packages. Correlation calculations were performed using custom R scripts. t-SNE and gap statistics calculations were performed as described by Grun et al. [41]

### Confocal microscopy

Images were obtained with a spectral confocal LEICA SP8 scanning microscope (Leica Microsystems, Germany). 5.10^4^ cells were cultured in a 48-well plate in 200μL prestimulation medium. After 72h, 100μL of 3% glutaraldehyde was added to the cell-containing well (1% final) for 15 minutes. Cells were washed twice with PBS 1X and incubated two hours with 2mg/mL NaBH4 at room temperature. Fcɤ receptors were saturated with Gamma Immune (Sigma Aldrich) for 5 min at 4°C (1:2 dilution). The cells were permeabilized with the fix/perm kit (BD-Biosciences), labeled for 20 minutes at 4°C with a 1:10 dilution of the mouse anti-humanCD133-APC antibody (clone Ac133 – Miltenyi Biotec), a 1:1000 dilution of phalloidin-TRITC (Sigma Aldrich) and stained with DAPI.

The images were acquired using a 63X PL APO CS2 1.40 NA oil immersion objective (Leica Microsystems, Germany). DAPI was excited with a 405nm, TRITC with a 552nm and APC with a 635nm laser. Finally, images were processed with a contrast enhancement algorithm (histogram equalization) and a home-designed background substraction algorithm.

### Micro-grid cell culture

A polydimethylsiloxane (PDMS) micro-grid array (Microsurfaces, Australia) of 1024 micro-wells (125 μm width, 60 μm-depth) was placed in specialized culture dish divided into 4 parts (Hi-Q4, Ibidi, Germany). Each part of the dish was filled with cell culture medium. A suspension of 5 × 10^3^ cells per case was added at a concentration likely to lead to a high number of wells with a single cell.

### Time-lapse microscopy

The time-lapse microscopy protocol was previously described [42]. Time-lapse acquisitions were performed with the Biostation IM time-lapse microscope (Nikon Instruments, Europe). 20 field positions were recorded covering 4 micro-wells each. Images were acquired every minute for 7 days using a 20X magnitude phase contrast objective. Only micro-wells containing a single cell were considered in the analyses.

### Image analyses

Images were analysed using ImageJ 1.47g 64-bits software (Rasband, W.S., ImageJ, U.S. National Institutes of Health, Bethesda, Maryland, USA, http://imagej.nih.gov/ij/, 1997–2014.). Cell tracking was performed manually using the ImageJ TrackMate plugin. The morphologies of first, second and third generation cells were analyzed semi automatically with Fiji (ImageJ 1.50e). A cell counter plugin was used to identify the moment when the cell switches from a round to a polarized morphology.

### Time-lapse data analyses

Analyses of time-lapse data were performed using R software. Cell lineage representations, cycle length, roundness and switch frequency were calculated with custom R made scripts. Euclidean distances of the last three parameters (cycle length, roundness and switch frequency) between the two sister cells were calculated. Cells were classified into 2 groups using the kmeans algorithm: derived from symmetric or asymmetric division. Boxplot representation combined with individual points was calculated with the beeswarm package (Aron Eklund (2016). beeswarm: The Bee Swarm Plot, an Alternative to Stripchart. R package version 0.2.3. https://CRAN.R-project.org/package=beeswarm).

### Proliferation assay

CD34+ cells were labeled with 2,5μM of CellTrace Violet (CTV) (Life technologies) at t=0h and analyzed using flow cytometry (LSRII – BD biosciences, France) after 24h, 48h and 72h with ModFit LT software as described previously by Neildez et al. [43]

### Image flow cytometry assay

Image flow cytometry analysis was performed using Image Stream MKII (Amnis, Proteigen, Merk Millipore). 5.10^4^ cells were cultured in a 48-wells plate in 200μL prestimulation medium. After 72h, 100μL of 3% glutaraldehyde was added to the cell-containing well (1% final) for 15 minutes. Glutaraldehyde offers good preservation of cell shape. Cells were washed twice with PBS 1X and incubated two hours with 2mg/mL NaBH4 at room temperature. Fcɤ receptors were saturated with Gamma Immune (Sigma Aldrich) for 5 min at 4°C (1:2 dilution). Cells were labeled for 20 minutes at 4°C with a 1:10 dilution of mouse anti-human CD133-APC antibody (clone AC133 – Miltenyi Biotec). Cell were then suspended in PBS and analysed with the image flow cytometer. Bright Field and APC channels were recorded (Bright Field: 745nm laser – APC: 642nm laser) with the 40X magnitude objective. Analyses of image stream data were performed with IDEAS® (Amnis, Proteigen, Merk Millipore).

### Cell sorting analysis

The CD34+CD133^high^ and CD34+CD133^low/neg^ cells were sorted at t=48h. Prior to labeling, Fc receptors were saturated with Gamma Immune (Sigma Aldrich). The CD34+ cells were labeled with CD34-PE (Miltenyi Biotec), CD45-APC-H7 (Beckman Coulter) and CD133-APC (clone AC133, Miltenyi Biotec) antibodies and 7-AAD marker (Sigma Aldrich). Isotype controls were used for the gating strategy. Cells were purified using a MoFlo® Astrios cell sorter (Beckman Coulter, France) and analysed with Kaluza software.

### Flow cytometric assay

The CD34+ cells were labeled using the following cell-surface markers: CD34-PE (Miltenyi Biotec), CD38-Pacific Blue (Beckman Coulter) and CD90-APC-Cy7 (Beckman Coulter) antibodies and 7-AAD marker (Sigma Aldrich). Isotype controls were used for gating strategies. Cells were analysed at 72h after prestimulation by flow cytomety (LSRII – BD biosciences, France) and analysed with FlowJo (v10.1) software.

## Author contribution

AP, DS, AM and OG designed the study

AM, AR, JC, CS and RP conducted the experiments

AM, DS, GC, JC and AP analysed the results and performed statistical analysis

AM, JC and DS prepared the figures

AP wrote the paper with the help of his colleagues

## Acknowledgements

This work was supported by EPHE (11REC/BIMO), ANR grant n° BSV6 014 02 « Stochagene » and Genethon. The authors are grateful to Jean-Jacques Kupiec and François Delhommeau for helpful discussions, to Peggy Sanatine for the precious help with cell sorting and Rhonda Campbell for the language revision of the manuscript. The authors declare no competing interest.

## Supplemental Information

**Table 1.**
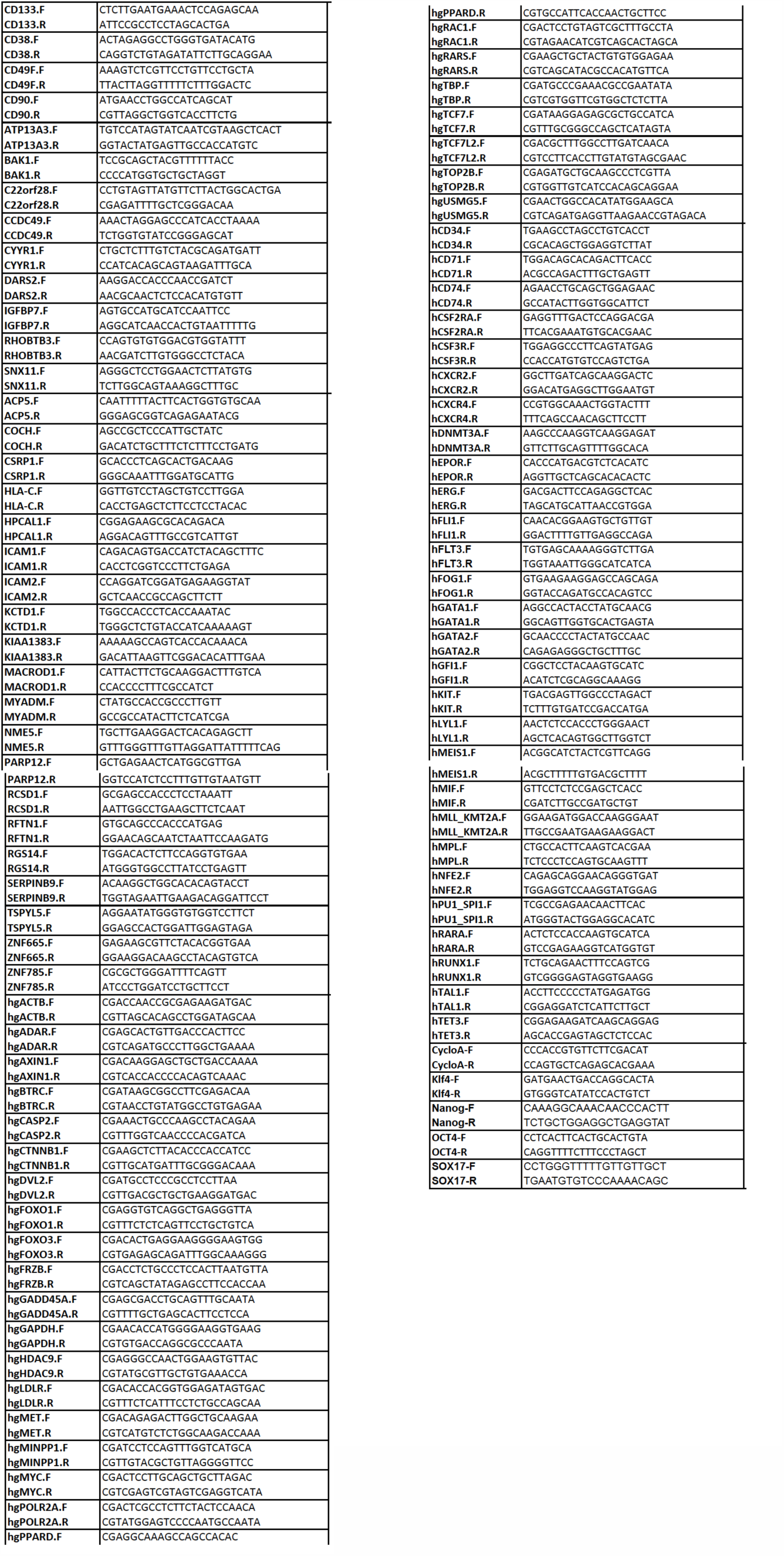
List of genes analyzed and primer sequences used for single-cell qRT-PCR amplification.

**Fig.S1.**
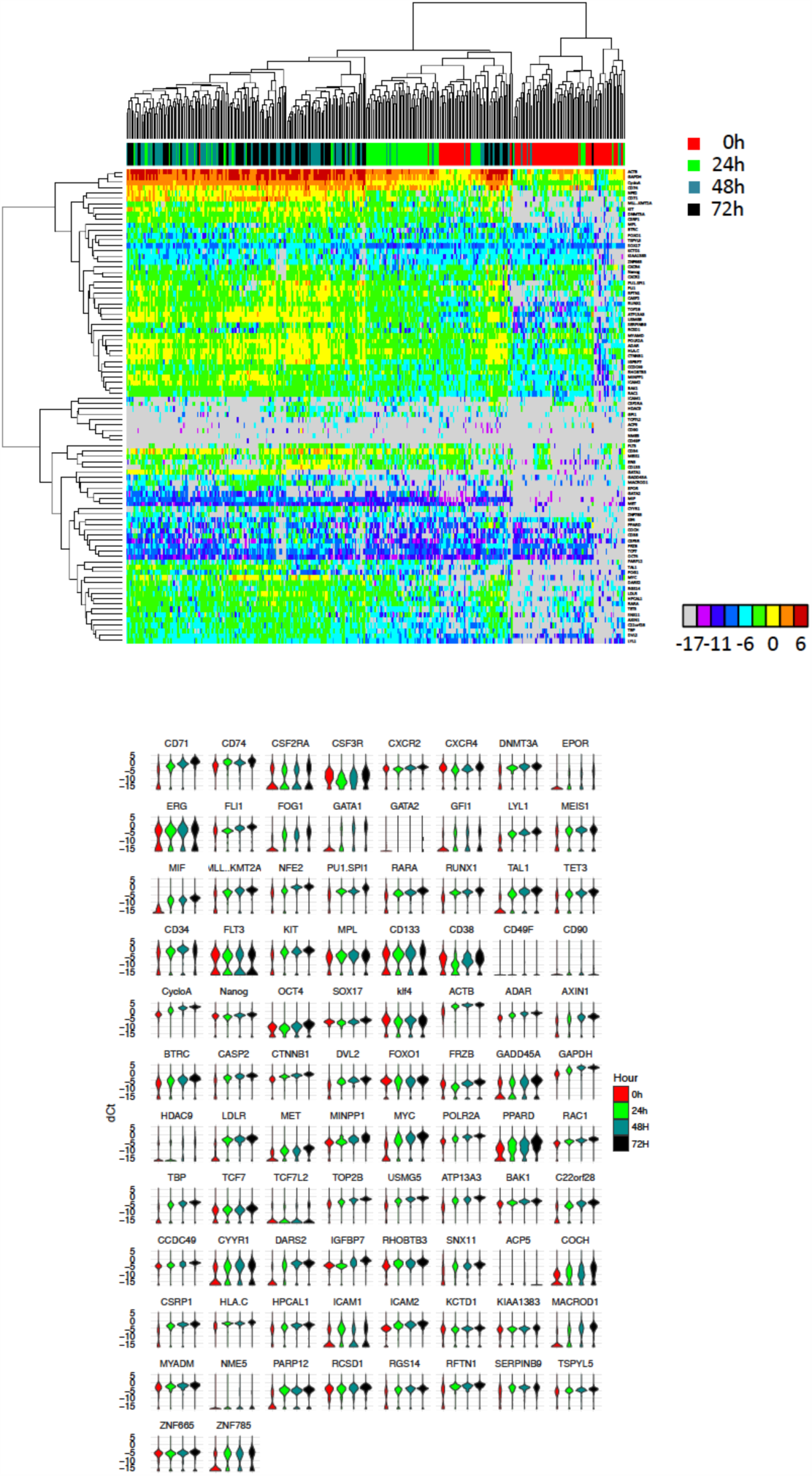
Full set of gene expression data obtained using single-cell qRT-PCR in cord-blood CD34+ cells cultured in vivo with early-acting cytokines. A. Extended heat map of the transcriptional profiles of cord blood-derived CD34+ cells at t=0h, t=24h, t=48h and t=72h after the beginning of the experiment. The color codes for the time points of cells are indicated on the right, the color code for expression level are indicated below the heat-map. Note the tendency of cells with the same time-points to cluster. B. Violin plot representation of individual gene expression levels at the four time-points.

**Fig.S2.**
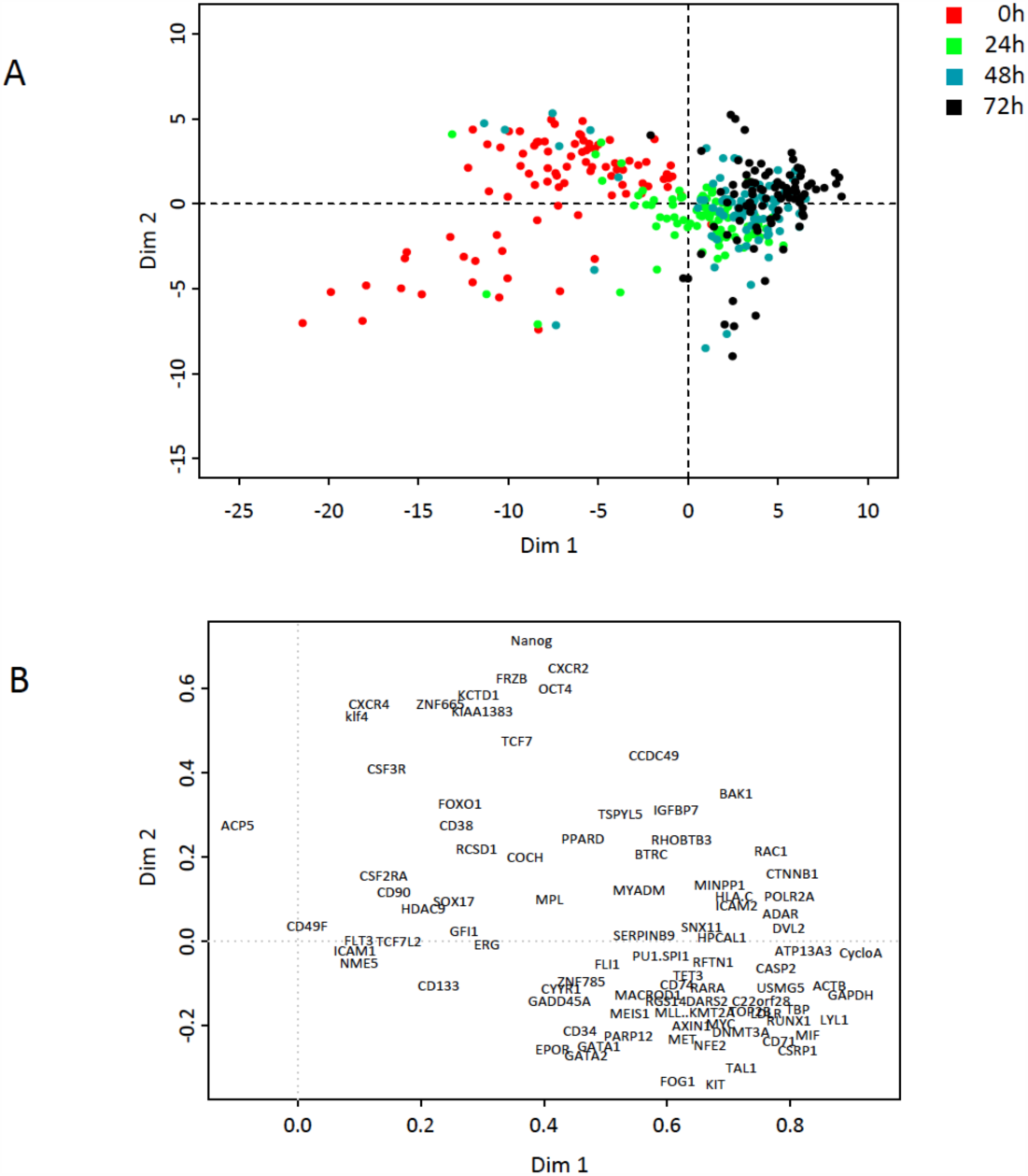
Principal component analysis of single-cell expression profiles. A. 2D PCA plot. Each point represents a single cell and the different time-points are coloured differently. Color codes are in the box to the right of the plot. B. Contribution of individual genes to principal component 1 and 2. Only the 40 highest contributions are indicated.

**Fig.S3.**
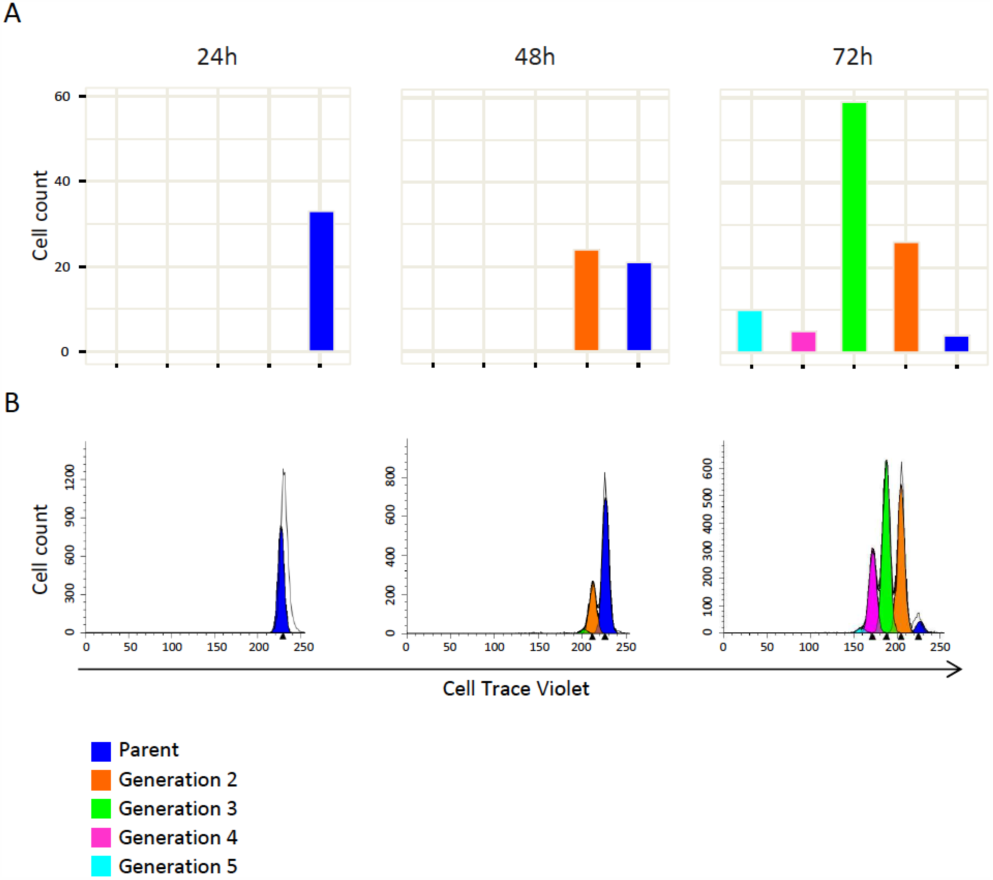
Analysis of cell division rates. A. The number of cells at t=24h, t=48h and t=72h as observed by time-lapse microscopy. The cells of different generations are color coded in the histogram. Note that none of the cells has divided after 24 hours and only 11 of the 32 cells underwent one division after 48 hours. At t=72h, three of the founder cells had not undergone division. B. Cell division analysis using Cell Trace Violet labelling. Cells were labelled at t=0h (not shown) and analyzed using flow cytometry at t=24h, t=48h and t=72h. When divided, the average fluorescence intensity of the two daughter cells is reduced by half compared to the maternal cell. Therefore, the peak on the right represents the parental generation. The number of the peaks to the left indicates the number of cell generations in the culture and the size of the peaks is indicative of the number of cells in each generation. Note that after 24h no cell division is detected and after 72h a fraction of undivided cells can still be detected. Most of the cells underwent one or two divisions. Overall, the profile is very similar to that detected by time lapse.

**Fig.S4.**
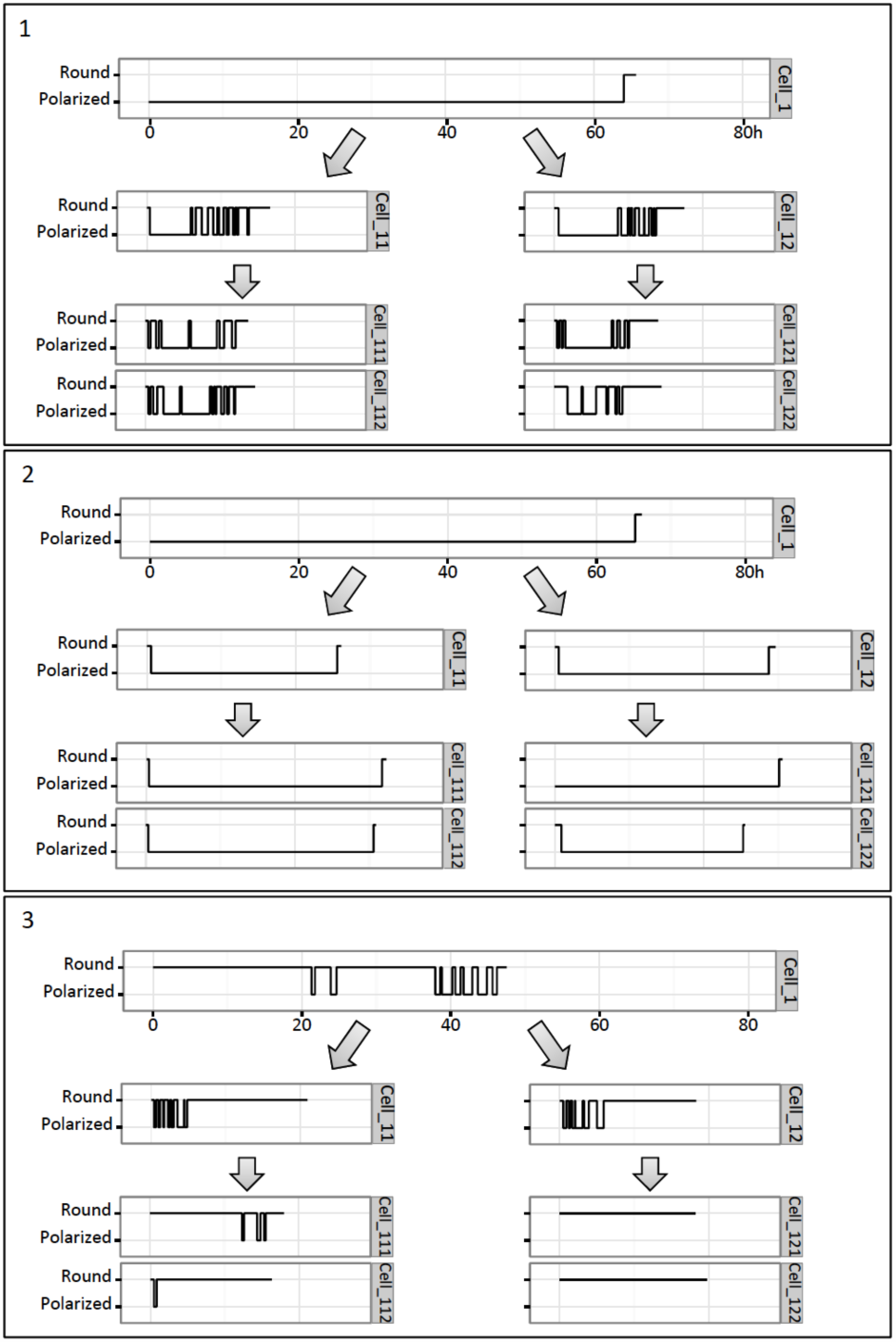
Representations of morphological profiles of cells in three representative clones. Each numbered panel represents a single clone. Each horizontal box in the three panels represents the morphology of an individual cell. The cell morphology – polarized or round – is shown with a horizontal line, the length of which is proportional to the time spent in the corresponding form. Vertical lines show the transitions between forms. The length of the horizontal lines is proportional to duration of the cell cycle and the time scale in hours is the same for each cell. The founder cell is numbered Cell_1, the two daughter cells Cell_11 and Cell_12 and granddaughter cell pairs as Cell_111, Cell_112 and Cell_121, Cell_122 respectively. In clone number 1 the polarized founder cell gives rise to frequent switcher daughters and granddaughters. Note the striking similarity of the time profiles for the morphological switches that can be observed in sister cells. In clone number 2 the polarized founder cell gives rise to stable polarized siblings. In clone number 3 the founder cell and its progeny are round. The two daughter cells switch to polarized shape for short periods. Note again the striking similarity of the sister cells’ switch profiles.

**Fig.S5.**
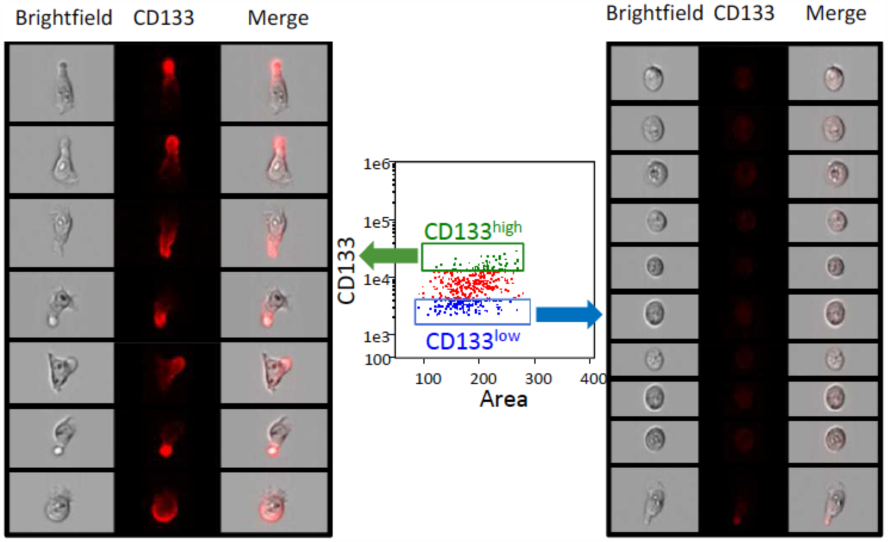
Cell morphology and CD133 localisation Image-based cytometry analysis shows correlation between the CD133 protein expression level and cell morphology at t=72h. The middle plot shows the CD133 protein density detected in glutaraldehyde-fixed cells. Representative examples of the morphologies of “high” (upper frame) and “low” (lower frame) expressing cells are shown on the left and right respectively.

**Fig.S6.**
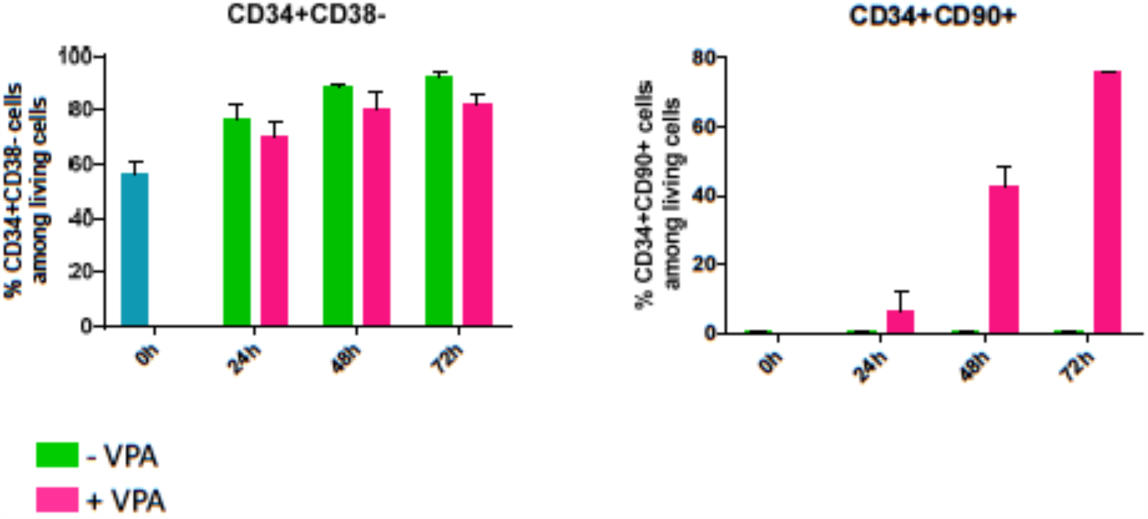
Cytometry analysis of the effects of valproic acid on CD34+ cells. The histogram in the left panel indicates the proportion of CD34+/CD38-cells in VPA+ and VPA- cell cultures at different time points. Note that there is no substantial difference between the two. The right panel indicates the proportion of CD34+/CD90+ cells in the same cultures. Note the increasing proportion of CD34+/CD90+ cells in VPA+ culture. This rapid increase cannot be explained by the selective proliferation of the CD90+ cells and is the result of the de novo synthesis of the CD90 protein, because as indicated in Fig.2, and Fig.S4, cells do not divide before 72h.

**Movie 1.** Time-lapse video of a cell clone with cells conserving polarized morphologies. The video has been accelerated to 5 frames per second.

**Movie 2.** Time-lapse video of a cell clone with cells conserving round morphologies. The video has been accelerated to 5 frames per second.

**Movie 3.** Time-lapse video of a cell clone with cells changing morphology at high frequency (dynamic phenotype of frequent switchers). Note that individual snapshots taken at different moments may show a population composed of only polarized, only round or cells with mixed morphology. The video is accelerated to 5 frames/sec.

